# ESMStabP: A Regression Model for Protein Thermostability Prediction

**DOI:** 10.1101/2025.02.18.638450

**Authors:** Marcus Ramos, Robert L. Jernigan, Mesih Kilinc

## Abstract

Accurately predicting protein thermostability is crucial for numerous applications in biotechnology, pharmaceuticals, and food science. Experimental methods for determining protein melting temperatures are often time-consuming and costly, driving the need for efficient computational alternatives. In this paper, we introduce ESMStabP, an enhanced regression model for predicting protein thermostability. To improve model performance and generalizability, we assembled a significantly larger dataset by combining and cleaning datasets previously utilized in other thermostability models. Building on DeepStabP, ESMStabP incorporates significant improvements, using embeddings from the ESM2 protein language model and thermophilic classifications. The predictions from ESMStabP outperform DeepStabP and other existing predictors, achieving an R^2^ of 0.95 and a Pearson correlation coefficient (PCC) of 0.97. Despite these improvements, challenges such as dataset availability. This work underscores the critical role of specific layer identification for model development and outlines potential directions for future advancements in protein stability predictions

## Introduction

Protein thermostability refers to a protein’s ability to maintain its structural and functional integrity over a range of temperatures (1). This characteristic is critical as proteins with low thermalstability are prone to denaturation and aggregation at higher temperatures, leading to loss of function or the formation of aggregates (2). Understanding and predicting the maximum temperature for protein thermostability is vital across various fields, including biotechnology, pharmaceuticals, and food science, where proteins are frequently exposed to temperature variations during cultivation, processing, and storage. The use of whole organisms in biotechnology make it necessary to identify which proteins are affected by increases in temperature. Obtaining accurate measures of protein thermostability for large numbers of proteins has been experimentally expensive. Experimental methods like differential scanning calorimetry (DSC) and circular dichroism (CD) spectroscopy are labor-intensive and time-consuming to obtain large numbers of protein melting temperatures (3). Computational-based prediction has emerged as an alternative, offering a more scalable and cost-effective solution. Current computational models often fall into two categories: those treating protein thermostability as a classification problem, predicting whether or not a given input sequence melts at a certain temperature (*T*_*m*_), and those treating it as a regression problem. Regression-based models, which predict the melting temperature from protein sequences, require a more nuanced understanding of thermostability. Recent advancements in deep learning, particularly in transformer-based protein language models, have been used to improve the accuracy of these predictions. ESM2 (4), a large language model (LLM) for protein sequences, captures intricate protein information by learning from vast amounts of sequence data. This model works by predicting masked letters in protein sequences, effectively learning the structural and functional aspects encoded within the sequences. Many current thermostability prediction methods leverage these language models since they capture extensive information about protein structure, and structure is hypothesized to be closely related to melting temperature. In this paper, we introduce ESMStabP, a regression model specifically designed to predict the melting temperature (*T*_*m*_) of proteins with enhanced accuracy. ESMStabP builds upon the foundation by another such model; DeepStabP (5), but incorporates several key improvements that set it apart. In the following sections, we will delve into the development and validation of ESMStabP, comparing its performance to DeepStabP and other models. We will also discuss the implications of these improvements for the broader field of protein thermostability prediction.

## Results

### Model Performance and Comparison

The results comparing ESMStabP to other protein thermostability prediction models are shown in Table 1. All models were trained on the same dataset to ensure a fair comparison. ESMStabP significantly improves protein thermostability prediction, with R^2^ and PCC values of 0.94 and 0.92, respectively, demonstrating a stronger linear relationship and better variance capture between predicted and actual melting temperatures. Despite this, ESMStabP shows slightly higher RMSE, MSE, and MAE values than DeepStabP when using the balanced dataset. When using the original, unbalanced dataset; ESMStabP performs better across all five metrics, achieving an MSE of 13.71, RMSE of 3.70, and MAE of 2.79, outperforming both DeepStabP and ProTstab2. Figure 1 illustrates the distribution of predictions made by ESMStabP, which follows a near-normal distribution slightly skewed below 0. While the model tends to slightly underestimate melting temperatures, its predictions remain relatively accurate, reflecting minimal bias.

**Table 1.**
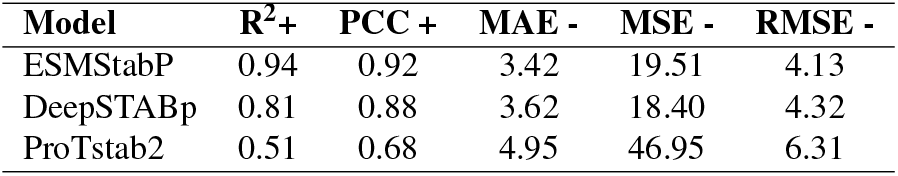
Comparison of other protein thermostability models for predicting a label *T*_*m*_. The “+” and “-” signs denote whether a higher or lower value is preferable for each metric

**Fig. 1.**
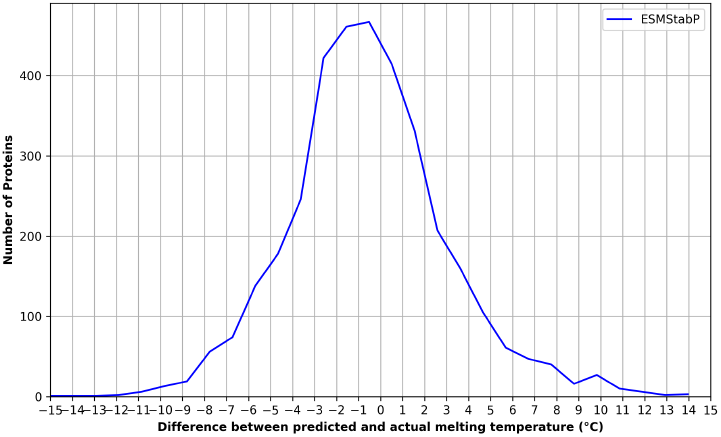
Distribution of the differences between predicted and label melting temperatures (*T*_*m*_) for ESMStabP. The overall spread of the distribution suggests that the model generally makes accurate predictions with minimal bias, as most predictions fall within a few degrees of the label values.

### Feature and Model Selection

In DeepStabP, optimal growth temperature (OGT) and indicating whether the experimental condition involved TPP studies with the heat treatment of cells or protein lysates were found to be effective features for predicting protein thermostability. Furthermore, TemBERTure (6) demonstrated that classifying sequences as either thermophilic or non-thermophilic prior to regression training led to improved results. In our approach, all three of these features—OGT, experimental condition (lysate or cell), and thermophilic classification—along with the output embedding from ESM2, are used as features for optimal results. While each feature was tested individually, incorporating all four features was found to yield superior results. Notably, excluding either OGT or the thermophilic class results in similar results; a PCC of 0.87 and 0.9 respectively, including both resulted in a significant improvement with a PCC of 0.97 despite both features being closely related. To identify the best-performing regression model, various models were experimented with, including linear regression, polynomial regression, and support vector regressor (SVR). Across all five evaluation metrics, the random forest regressor consistently performed the best.

### Layer Identification

Additionally, to evaluate the effectiveness of the individual layers within the ESM2 model, ESMStabP was tested with all 33 of the layer’s embeddings in isolation. Notably, a significant difference in performance was observed across the layers, with layer 33 demonstrating the best results, as shown in Figure 2. This outcome suggests that layer 33 incorporates the most relevant information for predicting protein thermostability. Consequently, for ESMStabP, only the embeddings from the 33rd layer were used, while the rest were ignored. Given the considerable variance in performance, this approach of layer isolation and identification may prove beneficial not only for other protein thermostability models but also for any models that leverage ESM2’s embeddings.

**Fig. 2.**
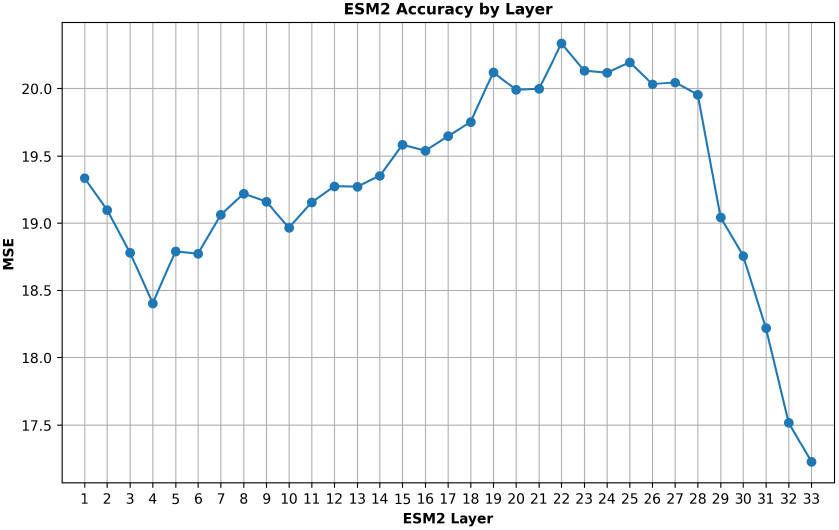
Mean squared error (MSE) for the regression models trained on each layer of esm2_t33_650M_UR50 for predicting the label melting temperature (*T*_*m*_).

## Materials and Methods

### A. Dataset Assembly

To ensure a fair comparison, the same dataset was utilized across all compared models. Protein sequences and their corresponding melting temperatures were included in this dataset, derived through high-throughput mass spectrometry-based thermo-proteome profiling (TPP) assays (7). In addition to the protein sequences and melting temperatures, information on whether the experimental condition was lysate or cell, as well as the optimal growth temperature for each protein, was contained in the dataset. The dataset was augmented further by classifiying whether or not the protein was thermophilic (growth temperature > 60°C) and non-thermophilic (growth temperature < 30°C) based on the success in doing so from TemBERTure; another protein thermostability prediction model. One problem with protein thermostability prediction is the lack of data due to how expensive it is to obtain the label melting temps. To solve this, we tracked down the datasets for many of the other thermostability models (5) (6) (8) and combined their data, removing duplicates and extracting the needed input features. The dataset was then separated into an 80% training and 20% testing split, and cross-validation was performed using K-fold (with K=5) to ensure robustness and reliability in the model evaluation. Finally, non-thermophilic proteins were randomly sampled to balance the dataset, as they were originally over represented.

### Extracting Protein Sequence Embeddings

All protein sequences from the dataset were then processed using the ESM2 protein language model, specifically the esm2_t33_650M_UR50 variant (available at https://github.com/facebookresearch/esm). Sequences were truncated to a maximum length of 1,022 residues to meet the model’s input requirements. The model’s output tensor logits were extracted for further processing. To produce a final embeddings tensor with a consistent dimensionality for regression models, the output tensor was averaged along the sequence length dimension. This resulted in a 3D matrix with dimensions 33 × 1,280, where 1,280 represents the embedding length of each residue, and 33 corresponds to the number of output layers in the esm2_t33_650M_UR50 model. Given the observed performance variance across layers, only the embeddings from the 33rd layer were utilized for further analysis, while the embeddings from The other layers were ignored. This decision was based on the finding that the 33rd layer contained the most relevant information for the target task. Each sequence’s embeddings were stored individually as numpy arrays in ‘.pt’ files.

#### Model Architecture

ESMStabP approaches protein thermostability prediction as a regression problem. Each of the columns from the augmented dataset, along with processed embedding output from esm2_t33_650M_UR50, were fed into a random forest regressor. The architecture of the model is depicted in Figure 3. Similar to DeepStabP, ESMStabP is easily extendable with other language models and input features. When experimenting with the components of DeepStabP, it was found that using the inputs as raw features was more effective than utilizing multi-layer perceptrons. The final model is available in joblib format at https://github.com/marcusramos2024/ESMStabP/tree/main

**Fig. 3.**
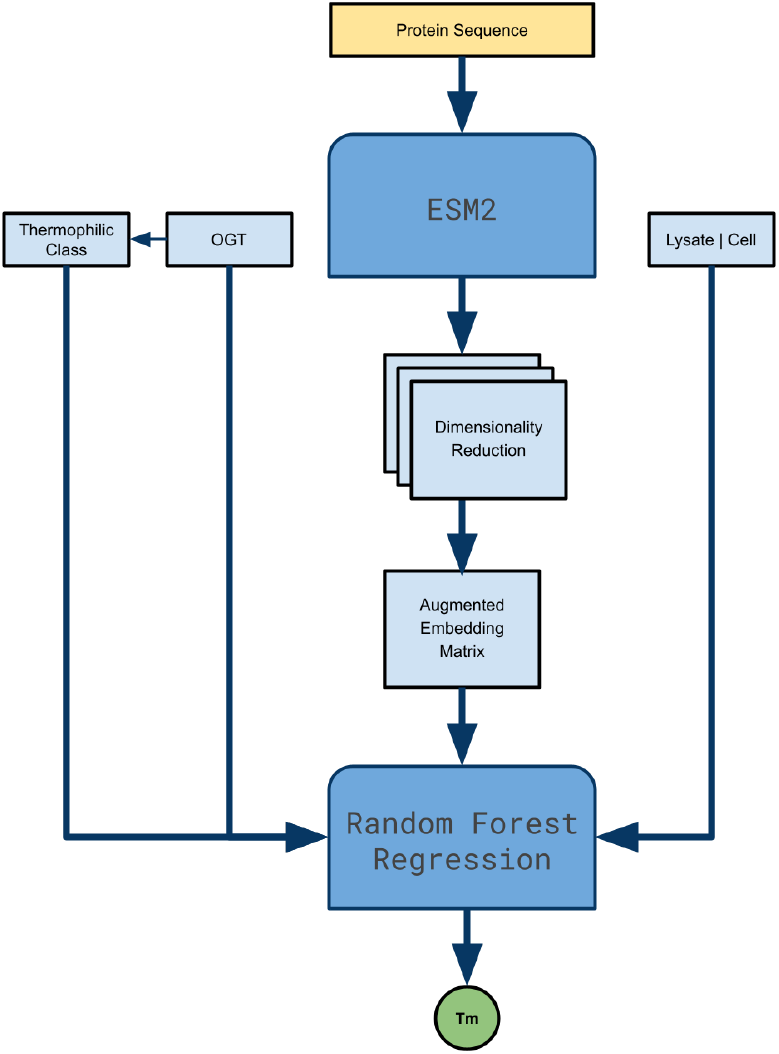
Visual representation of ESMStabP’s end-to-end architecture, highlighting Thermophilic class, Optimal Growth Temperature (OGT), and Experimental Condition as input features.

#### Evaluation Metrics

To validate the performance of the models during training and testing and to enable fair comparisons with alternative approaches, several of the most commonly used evaluation metrics were computed. Each metric measures the discrepancy between vectors of *N* experimentally determined *T*_*m*_ (*y*) and predicted *T*_*m*_ (*ŷ*).

Coefficient of Determination (*R*^2^):

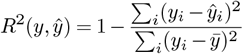

Mean Average Error (MAE):

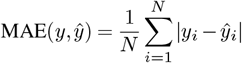

Mean Squared Error (MSE):

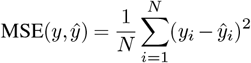

Root Mean Squared Error (RMSE):

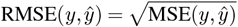

Pearson Correlation Coefficient (PCC):

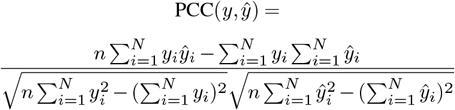

## Discussion

### Previous Models

Advancements in protein thermostability prediction have been driven by models that use embeddings from protein language models as input features, leading to improved accuracy and applicability. An earlier model, ProTstab2 (9), was notably successful in training on these embeddings. This approach allowed for more nuanced predictions by capturing the complex structural information embedded within protein sequences. However, ProTstab2 faced limitations in terms of accuracy and generalizability. DeepStabP improved upon ProTstab2 by integrating more advanced techniques and features. It utilized embeddings from the ProtTrans-XL protein language model (10), which provided richer and more detailed representations of protein sequences. Additionally, DeepStabP incorporated multilayer perceptron (MLP) outputs to clarify experimental conditions—such as whether the study involved lysate or cell samples and included the optimal growth temperature. These enhancements led to better performance in predicting protein thermostability, demonstrating the importance of incorporating experimental conditions and growth temperature into the model. However, DeepStabP was limited by its training data. The dataset used was biased; with significantly more nonthermophilic sequences than thermophilic ones. Consequently, TemBERTure (6) introduced an approach with a balanced dataset. TemBERTure also found succsess through incorporating a thermophilic classification layer and utilizing the BERT (11) protein language model.

### Parameter-Efficient Fine-Tuning with LoRA

Recent investigations into parameter-efficient fine-tuning methods for large protein language models have shown encouraging results when applied to specific problems. In particular, a study (12) demonstrated that Low-Rank Adaptation (LoRA) can be successfully applied to protein language models to specialize them for various prediction tasks without requiring full model retraining. Inspired by these findings, we implemented LoRA-based parameter-efficient fine-tuning on two of the leading protein language models; ESM2 and Prot5. Although ESM2 consistently outperformed Prot5, our experiments revealed that simply fine-tuning these models with LoRA did not surpass the predictive accuracy of ESMStabP. This suggests that the information captured within the language models is not sufficient on its own and that feature identification remains a critical factor in achieving high predictive accuracy.

### Improvements and Limitations

Our research highlights the Random Forest Regressor’s superior performance over Support Vector Regressor (SVR) and both linear and logistic regressors traditionally used in thermostability models. Additionally, using raw values as input features proved more effective than employing multi-layer perceptrons (MLPs). The model’s performance was further optimized by incorporating input features from both DeepStabP (e.g., Optimal Growth Temperature and experimental conditions) and TemBERTure (e.g., thermophilic classification), along with the more capable ESM2 model, replacing ProtTrans. One notable contribution is the identification of layer isolation’s importance within the ESM2 model, yielding substantial improvements for similar problems that utilize large language model embeddings. ESMStabP’s flexibility is also enhanced by its ability to function effectively without the need for specific experimental conditions or growth temperature data, unlike DeepStabP. This flexibility broadens its applicability across different research scenarios where such data might be unavailable or incomplete. Finally, by contributing a singular unified, thoroughly curated dataset, we establish a valuable resource that can be readily leveraged for future models and thermostability research questions. Despite these improvements, ESMStabP has limitations. Even with our larger combined dataset, the sourced datasets were inherently biased, with non-thermophilic proteins being more abundantly represented. After balancing the dataset to address this bias, the size of the dataset was significantly reduced, which could impact the model’s training and performance. This issue underscores the necessity for greater diversity in datasets to enhance the reliability and generalizability of protein thermostability models. Additionally, the dataset used in this study lacks a wide variety of organisms, limiting the model’s generalizability across different species. Future work should focus on expanding the dataset to include a more diverse array of proteins from various organisms, thereby enhancing the model’s robustness and applicability. While this study tested only a few models (ESM2, ProtTrans-XL, and BERT), there are many other models that could be explored in future research to further improve thermostability prediction.

## Acknowledgement

This work was supported by DOE grant DE-SC0022090 and NIH grant R01HG012117.

## Bibliography

1. Cathal Ó Fágáin. Understanding and increasing protein stability. Biochimica et Biophysica Acta, 1252:1–14, 1995. doi: 10.1016/0304-4157(95)00003-4.

2. John H. Bischof and Xiaoming He. Thermal Stability of Proteins. Annals of the New York Academy of Sciences, 1066:12–33, 2006. doi: 10.1196/annals.1363.003.

3. Joachim Seelig and Helmut-J. Schönfeld. Thermal protein unfolding by differential scanning calorimetry and circular dichroism spectroscopy: Two-state model versus sequential unfolding. Quarterly Reviews of Biophysics, 49:e9, 2016. doi: 10.1017/S0033583516000044.

4. Alex Rives, Joshua Meier, Tom Sercu, Siddharth Goyal, Zeming Lin, Jason Liu, Demi Guo, Myle Ott, C. Lawrence Zitnick, Jianyuan Ma, and Rob Fergus. Biological structure and function emerge from scaling unsupervised learning to 250 million protein sequences. Proceedings of the National Academy of Sciences, 118(15), 2021. doi: 10.1073/pnas.2016239118.

5. Florian Jung, Karl Frey, Daniel Zimmer, and Tobias Mühlhaus. DeepSTABp: A deep learning approach for the prediction of thermal protein stability. International Journal of Molecular Sciences, 24(8):7444, 2023. doi: 10.3390/ijms24087444.

6. Chiara Rodella, Symela Lazaridi, and Thomas Lemmin. TemBERTure: advancing protein thermostability prediction with deep learning and attention mechanisms. Bioinformatics Advances, 4(1):vbae103, 2024. doi: 10.1093/bioadv/vbae103.

7. André Mateus Nina Kurzawa, Ingo Becher, Sudhakar Sridharan, Daniel Helm, Fabian Stein, Athanasios Typas, and Mikhail M. Savitski. Thermal proteome profiling for interrogating protein interactions. Molecular Systems Biology, 16:e9232, 2020. doi: 10.15252/msb.20199232.

8. Mengyu Li and et al. DeepTM: A deep learning algorithm for prediction of melting temperature of thermophilic proteins directly from sequences. Computational and Structural Biotechnology Journal, 21:5544–5560, 2023.

9. Y. Yang, J. Zhao, L. Zeng, and M. Vihinen. ProTstab2 for Prediction of Protein Thermal Stabilities. International Journal of Molecular Sciences, 23:10798, 2022. doi: 10.3390/ijms231810798.

10. A. Elnaggar, M. Heinzinger, C. Dallago, G. Rehawi, Y. Wang, L. Jones, T. Gibbs, T. Feher, C. Angerer, M. Steinegger, D. Bhowmik, and B. Rost. ProtTrans: Toward Understanding the Language of Life Through Self-Supervised Learning. IEEE Transactions on Pattern Analysis and Machine Intelligence, 44:7112–7127, 2022. doi: 10.1109/TPAMI.2021.3095381.

11. J. Devlin, M. Chang, K. Lee, and K. Toutanova. BERT: Pre-training of Deep Bidirectional Transformers for Language Understanding. arXiv (Cornell University), 2018. doi: 10.48550/arxiv.1810.04805.

12. Robert Schmirler, Michael Heinzinger, and Burkhard Rost. Fine-tuning protein language models boosts predictions across diverse tasks. Nature Communications, 15:7407, 2024. doi: 10.1038/s41467-024-51844-2.

